# Cryo-ET reveals two major tubulin-based cytoskeleton structures in *Toxoplasma gondii*

**DOI:** 10.1101/2021.05.23.445366

**Authors:** Stella Y. Sun, Li-av Segev-Zarko, Muyuan Chen, Grigore D. Pintilie, Michael F. Schmid, Steven J. Ludtke, John C. Boothroyd, Wah Chiu

**Author notes:** These authors contributed equally (Stella Y. Sun, Li-av Segev-Zarko, Muyuan Chen). Correspondence (J.C.B), (W.C.).

## Abstract

In the obligate intracellular parasite, *Toxoplasma gondii*, the subpellicular microtubules (SPMTs) help maintain shape, while the apical conoid (also tubulin-based) is implicated in invasion. Here, we use cryo-electron tomography to determine the molecular structures of the SPMTs and the conoid-fibrils (CFs) in vitrified and detergent-lysed parasites. Subvolume densities from detergent-extracted parasites yielded averaged density maps at subnanometer resolutions, and these were related back to their architecture *in situ*. An intraluminal spiral (IS) lines the interior of the 13-protofilament SPMTs, revealing a preferred orientation of these microtubules relative to the parasite’s long axis. Each CF is composed of 9 tubulin protofilaments, that produce a comma-shaped cross-section, plus additional associated components. Conoid protrusion, a crucial step in invasion, is associated with an altered pitch of each CF. The use of basic building blocks of protofilaments and different accessory proteins in one organism, illustrates the versatility of these critical structures.

## INTRODUCTION

*Toxoplasma gondii* belongs to the phylum Apicomplexa which consists of a diverse group of obligate intracellular parasites. This single-celled eukaryote possesses a broad host range, capable of infecting almost any nucleated cell in any warm-blooded animal (Dubey, 2004). Symptoms associated with *Toxoplasma* infection are diverse, ranging from mild to fatal, especially in the developing fetus or immunocompromised patients (e.g., those with AIDS, heart transplants, etc.). Due to its prevalence worldwide, *Toxoplasma* presents a global health problem.

Upon infection, *Toxoplasma* tachyzoites proliferate rapidly within the host cell and eventually cause host cell lysis. Released free tachyzoites are highly motile, actively invading more host cells. To accomplish this, they deploy a sophisticated and complex strategy involving a specialized, apical architecture and tightly choreographed secretion from the apical organelles including micronemes and rhoptries (Ben Chaabene et al., 2020; Carruthers and Boothroyd, 2007; Dubois and Soldati-Favre, 2019). Among the events associated with invasion is remodeling of the cytoskeleton at the apical end, including protrusion of the conoid, a prominent bundle of spirally organized fibers (Graindorge et al., 2016; Hu et al., 2002; Leung et al., 2020; Lycke et al., 1975).

The tubulin-based cytoskeleton that forms the pellicle and apical complex of *Toxoplasma* is not only responsible for cellular structural integrity, but also provides a mechanosensitive scaffold for cell motility and a polarized discharge of key invasion factors (Dos Santos Pacheco et al., 2020; Morrissette and Gubbels, 2020). These crucial roles make these tubulin-based structures attractive drug targets for treating infection. Previous proteomic analysis of tachyzoite cytoskeletons identified both α- and β-tubulin units to be present (Hu et al., 2006; Nagel and Boothroyd, 1988), and a novel tubulin polymer in the conoid has been observed in detergent-extracted cytoskeleton (Hu et al., 2002). The precise molecular organization of tubulins in the apical complex remained unclear, making it difficult to study how they interact and function during infection.

Cryo-electron tomography (cryo-ET) is a powerful tool to study molecular structure *in situ* (Chen et al., 2017b, 2019; Robinson et al., 2007; Tegunov et al., 2021). This method can define the spatial context of cellular architecture without using chemical fixatives or metal stains. *Toxoplasma* tachyzoites are crescent-shaped and approximately 2 by 6 μm. Although these cells are generally too thick for studying intact parasites by cryo-ET, our interest is in the tapered, apical region, which is about 400 nm thick, thin enough for a 200-300 kV electron beam to form images. In this study, we focused on detailed image analysis of the SPMTs and CFs in intact parasites as well as in detergent-extracted cytoskeletons using subtomogram averaging. Our analysis revealed unexpected structural differences in the tubulin protofilament organization of SPMTs and CFs at subnanometer resolution and enabled detection of their specific association with additional, yet-to-be-identified molecular components. Dissection of the components in apical complexes containing SPMTs and CFs provides a detailed understanding of how different tubulin-based macromolecular complexes are assembled in a cellular context and provides clues to their crucial roles in parasite biology.

## RESULTS

### Cryo-ET reveals novel structural details in the apical complex

CFs and SPMTs are two types of tubulin-based cytoskeletal elements that maintain the shape and polarity of the apical complex, a key part of the invasion machinery in *Toxoplasma*. Under calcium flux, a chemical stimulus critical to the invasion process in this phylum of parasites, the CFs protrude from the surrounding SPMTs (Mondragon and Frixione, 1996), making the apical end of the parasites fall well within the thickness limitation of cryo-ET imaging (~500nm). Using phase optical microscopy, we assessed the percentage of CF protrusion in extracellular parasites following calcium ionophore (A23187) treatment. Examination of tachyzoites before loading them on the EM grid showed ~87% presented a protruded conoid, ensuring a large number of parasites suitable for imaging. To maximize the recognition of distinct subcellular components in this structurally complex area of the cell, we used a Volta phase plate (VPP) located in the focal plane of the objective lens of the electron microscope (Danev et al., 2014). The resulting enhancement of image contrast facilitates the use of convolutional neural networks (CNN) (Chen et al., 2017b) for annotation of distinct secretory organelles and cytoskeleton elements in the apical complex. Figure 1 and Movie S1 show data for a representative tomogram generated in this way. Among the features revealed were the tubulin-based conoid-fibrils (CFs) and intraconoidal microtubules (IMTs), bounded by two distinct preconoidal rings (PCRs) on top and the apical polar ring (APR) on the bottom, with micronemes, rhoptries and other organelles filling its interior space (Figure 1B, E). The SPMTs appeared to be aligned to the inner surface of the pellicle along the entire longitudinal axis of the cell body that appears in our field of view. For tachyzoites presenting intermediate or completely retracted conoids, we can capture the retracted conoid shielded by the ring of SPMTs.

**Figure 1.**
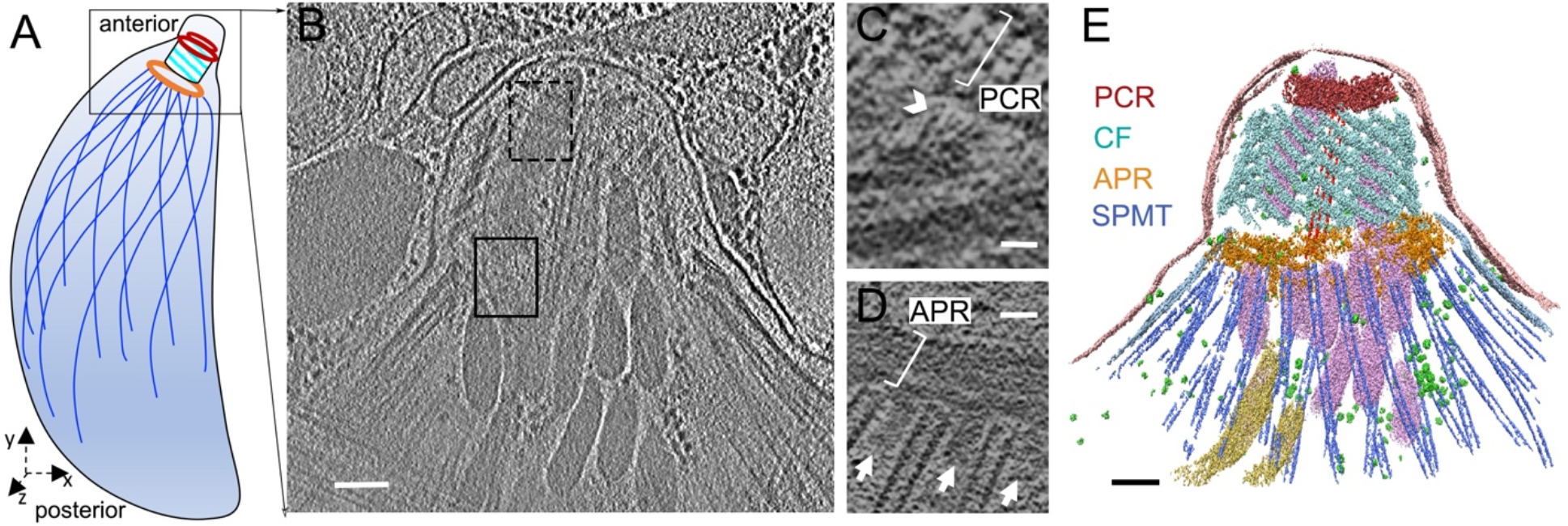
Three-dimensional organization of the apical complex in *Toxoplasma* tachyzoites reveals the subpellicular microtubule and conoid architecture (A) Cartoon of the *Toxoplasma* tachyzoite, the life stage which has been imaged by cryo-ET in (B) and annotated in (E). (B) Tomographic slice of a representative apical complex of an intact tachyzoite recorded with Volta phase plate optics. (C) A zoomed in view of a tomographic slice in the dashed black box of (B), showing two CFs in the xy plane; their anterior ends extending towards a preconoidal ring (PCR; white bracket) with filamentous density (a white arrowhead) between them. Scale bar, 25nm. (D) Portion of a tomographic slice in the black box of (B), showing the SPMTs in the xy plane, and their association with the apical polar ring (APR; white bracket). Note columnar densities (white arrows) emerging from the APR and positioned between the SPMTs. Scale bar, 25nm. (E) 3D segmentation of the tomogram shown in (B) including the PCR (red), conoid fibrils (CFs, cyan), APR (golden) and SPMTs (blue), intra-conoidal microtubules (red), micronemes (pink), rhoptries (yellow) and plasma membrane (pale pink). Scale bar, 100nm. The black squares represent the general area that is expanded in (D), using tomographic slices optimal for the features being discussed. The dashed square indicates that the slice in (C) is taken from the back side of the conoid, with the CFs extending from lower left toward upper right, opposite from the front view. Scale bar, 100nm. A movie with the complete tomogram is available in Movie S1.

Transverse slices of the CF from the reconstructed tomograms exhibit a comma shape, consistent with previous negative staining electron microscopy studies (Hu et al., 2002). Close examination of the apical end of the CF revealed filamentous densities extending toward the PCR both in intact parasites and in detergent-extracted cytoskeleton, showing a close connection between the conoid and the PCR (Figure 1C and Figure S1A-D). These feature discoveries were enabled by the use of the VPP imaging. We aligned and averaged 50 subvolumes of the region that includes both the PCR pairing units, the anterior end of the CF and the filamentous densities from 6 cellular tomograms. The averaged map shows that one CF spans about 4 ring-pairing units of the PCR (Figure S1E and F). None of the filaments which are visible in the tomogram was revealed in this averaging processing probably because they are not oriented in the same way. We annotated the reconstructed tomogram to directly examine the number of assembled CFs per apical complex but the complexity and image limitations hindered our ability to trace them reliably. To maximize the visibility of the individual CF, we annotated the CFs in the deoxycholate-treated parasites, whose conoid area is thin enough to be fully annotated; the results showed that each conoid consists of 14 (n=6) or 15 (n=4) CFs (Figure S2A-D).

The SPMTs span most of the cell body length, contributing to the elongated shape of toxoplasma tachyzoites due to their tight connection to the inner membrane complex (Morrissette et al., 1997). The SPMTs are defined as polar structures, emanating from the APR. Examination of the intact cell tomograms in the area of the APR revealed microtubules that are regularly and radially spaced around the ring with additional, discrete density appearing between neighboring SPMTs (Figure 1D). Annotation of the 3D volume allows us to count the number of SPMT per cell that otherwise overlap or would be completely missed in a 2D slice view. To avoid potential uncertainty from intraconoidal or broken microtubules, we counted only the SPMTs emanating from the APR of intact parasites. The results showed that 15 of our tomograms displayed the previously reported number of 22 SPMTs, while 10, 6 and 1 tomograms showed 21, 23 and 24 SPMTs per cell, respectively (Figure S2E, F).

### SPMTs contain intraluminal spirals of unknown components, with unique organization at the seam

To investigate the structural details of the SPMTs, detergent-extracted cells were vitrified, imaged and examined. The extracted cells are free of loosely bound cellular contents, thereby maximizing the signal-to-noise ratio of the remaining highly organized cytoskeleton components. Restricted by the field of view of the electron microscope camera, the imaging magnification was chosen to optimally cover the microtubules as they emerged from the APR. The SPMTs remain connected to the APR, together with the unknown, interspersed pillar densities that were detected in the tomograms of intact parasites (Figure 1D and 2A). To obtain an intermediate resolution structure of these SPMTs, we performed subvolume alignment and averaged with 8nm periodicity, based on the expected repeat for tubulin heterodimers and as also observed in the intraluminal spiral (IS) density repeats in Figure 2A and previous reports (Cyrklaff et al., 2007). It has been shown previously that SPMT has a standard microtubule arrangement (Hu et al., 2002; Nichols and Chiappino, 1987); our subvolume averaging assumed the tubulin protomers followed the microtubule-specific pseudo-helical symmetry (Sui and Downing, 2010), in which each microtubule consists of 13 protofilaments of alternating α- and β-tubulin, with a unique junction called the “seam”, where α- and β-tubulin interact laterally. This interaction is different from the other interactions between the protofilaments. Beginning and ending at the seam, there is a 12 nm axial spacing of a single spiral of α- or β- subunits. Our subvolume averaging resulted in a 6.7 Å resolution map for a SPMT segment over a length of 24nm (Figure 2C and E and Figure S6C). Visualizing the density, we were able to identify 5-7 Å-wide rod-shaped features, characteristic of α helices, corroborating the subnanometer resolution estimate. The dispositions of these helices are consistent with the atomic structure of the tubulin molecules (Zhang and Nogales, 2015). However, the smoothness of the helix density suggests that the resolution appears to correspond to structural features expected in 7-8 Å resolution cryo-EM maps of single particles (Chiu et al., 2005). Though our high-resolution results of individual monomers of tubulin or IS are derived from this map with the microtubule pseudo-helical symmetry imposed, we also created a subtomogram average without imposing any symmetry (c1) using a search template having a well-defined seam, hereby called “symmetry-released” (see Methods). This was used for most of the interpretation of the topological organization of the SPMT and its associated IS.

**Figure 2.**
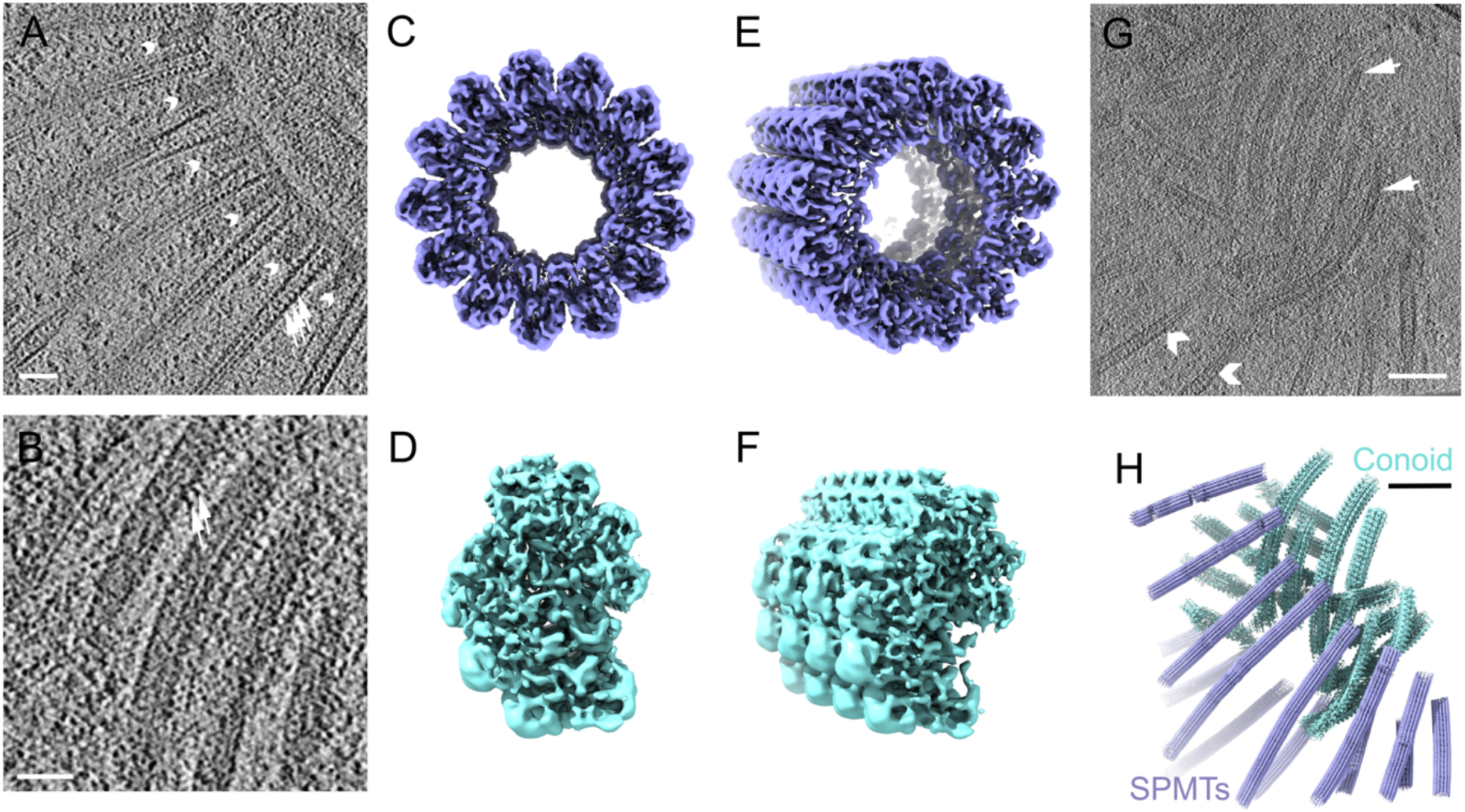
Determination of the 3D structures of apical SPMT (blue) and CF (cyan) segments in detergent extracted cytoskeleton of *Toxoplasma* (A) Portion of a tomographic slice showing the SPMTs with internal striation density (white arrows) and short, interspersed pillars (white arrowheads). Scale bar 50nm. (B) Portion of a tomographic slice showing individual CFs. Scale bar 50nm. (C, E) Reconstruction of a representative SPMT segment viewed from top and a tilted angle. (D, F) The top view and tilted view of a reconstructed CF segment showing an asymmetric, semi-circular profile. (G) A representative tomographic slice of the apical end cell extract, containing individual SPMTs (white arrowheads) and conoid fibrils (white arrows). Scale bar 100nm. (H) The three-dimensional organization of extracted SPMTs and CFs in the local region of the reconstructed tomogram shown in (G) based on the refined coordinates of individual particles.

It is known that *Toxoplasma*’s α- and β-tubulin sequences are ~85% identical to mammalian tubulins (Nagel and Boothroyd, 1988). To distinguish the α- from the β-tubulin isoforms in the averaged density map, we generated a homology model of *Toxoplasma* tubulins. While the *Toxoplasma* genome carries three α-tubulin genes, we used the protein sequence of α-1 (TGME49_316400) for the homology model as it is the only one detected in a proteomics study of the tachyzoite cytoskeleton (Gómez de León et al., 2014; Hu et al., 2006). The *Toxoplasma* genome also carries three β tubulin genes (TGME49_266960 221620 and 212240) with all isoforms reportedly detected in the same proteomics study, although transcriptomic studies indicated the β-3 isoform (TGME49_212240) is essentially not expressed while the other two are abundantly expressed in tachyzoites. By amino acid sequence, β-1 (TGME49_266960) and β-2 (TGME49_221620) share 96.9% identity and 98.9% similarity; given this near complete identity, we chose the β-1 isotype at random for modeling purposes. The α-1 tubulin model displays a longer S loop compared to β-1, corresponding to residues 359-372 in the α-1 protein and similar to the resolved structure of the porcine microtubule. Subtomogram averaging resulted in a map where the α-tubulin is readily differentiated from the β-tubulin unit due to the difference in the S loop (Figure 3B and C). Based on the correlation and visual analysis between the segmented tubulin subunit density in the subtomogram average and the map corresponding to ~8 Å resolution computed from the porcine α- and β-tubulin model (EMD-6439), we were able to distinguish these two subunits in our density map (Figure. 3A).

**Figure 3.**
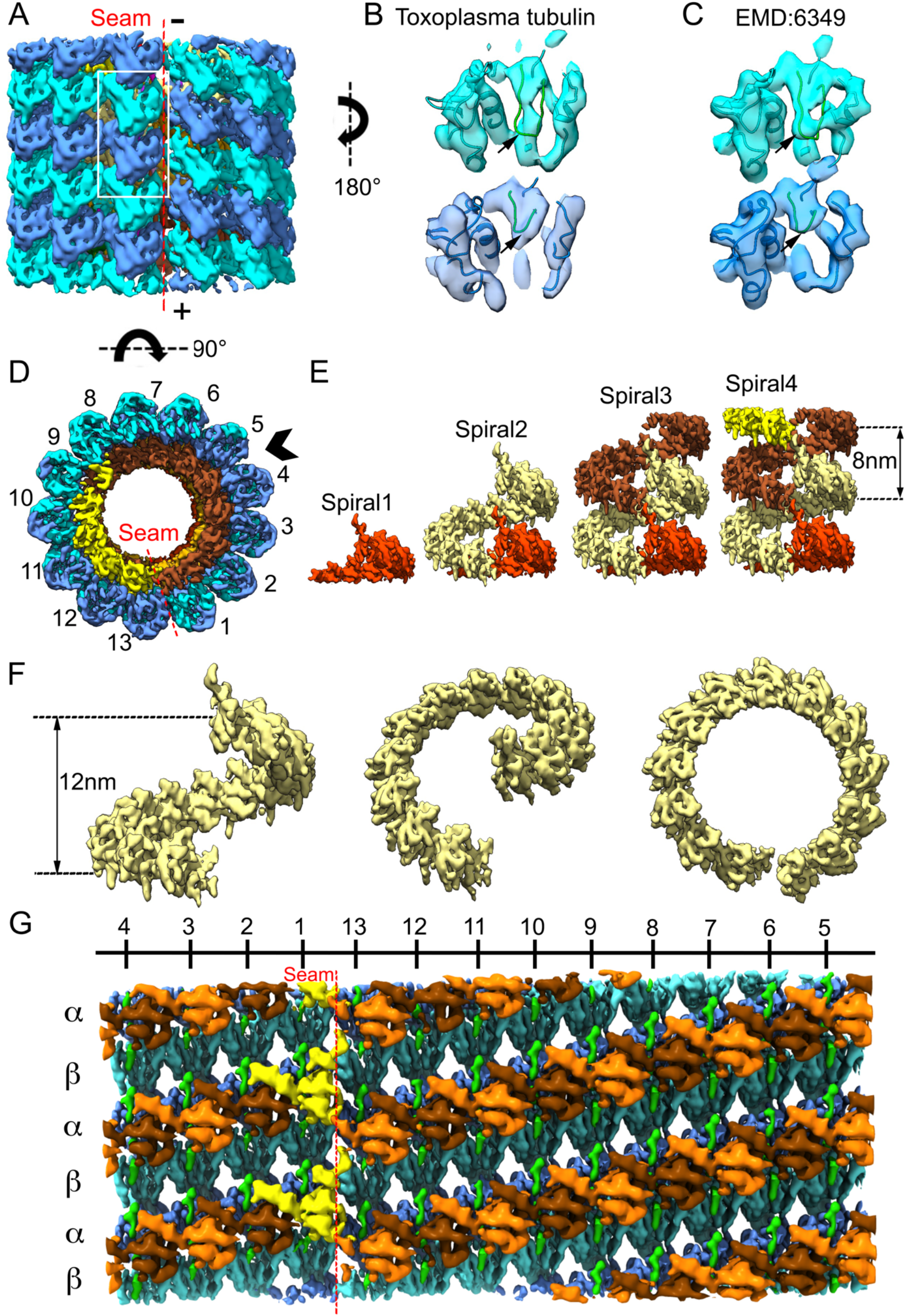
Structural analysis of SPMT tubulins and their associated lumen complex (A) The arrangement of α- (cyan) and β- (blue) tubulin subunits in the averaged density map of the SPMT segment, viewed from the seam, and showing the arrangement of the 13 protofilaments. The plus (+) and minus (−) ends of the microtubule are as indicated. The minus (up) end is toward the APR. (B) Density for an αβ-tubulin dimer as represented by the white boxed area of (A), viewed from within the SPMT lumen. Note the S-loop (green loop indicated by black arrows) in α-tubulin has extended density, but not in β-tubulin; this is validated by (C), an αβ-tubulin dimer segmented from a porcine microtubule (EMD6349) showing the S loop density of differences displayed at 8Å, also viewed from the lumen side. (D) Non-tubulin densities inside the tubulin-spiral, viewed within a cross-section of SPMT, are colored sienna and yellow. (E) Stacked spiral architecture of the IS complex with 4 stacked spirals shown in red, khaki, sienna and yellow. (F) One stack of the IS viewed from the seam at different angles. (G) Unwrapping the SPMT between protofilaments 4 and 5 shows the SPMT assembly from the inside, with an intraluminal spiral (IS) density axially arrayed along the tubulin lattice with 13 apparent repeating units. Unit X_1_ over the seam is highlighted in yellow; X_2_-X_13_ is shown in alternating sienna and orange; the rod-shaped densities between tubulin and X_1_-X_13_ units are shown in green.

In addition to the tubulin protein densities in the SPMTs, we observed additional densities (herein called the intraluminal spiral, IS) clearly visible in the cross-sectional view, rotating around the lumenal surface of the microtubule protofilaments (Figure 3D and E). Viewing from the longitudinal axis of the SPMT segment, this IS complex follows a left-handed and discontinuous helical pattern, built up stack by stack, similar to a series of split-ring washers (Figure 3F-G). The axial repeat of the washers is 8nm along the microtubule axis and the two split ends of each washer are 12 nm apart (Figure 3F), similar to the spirals of α- or β-tubulin. Based on the start and stop ends of each washer, we can reliably segment 13 structurally similar repeating units X_1_-X_13_, each estimated to be ~25kDa based on the volume density (Figure S3) (Pintilie et al., 2010). In examining the density features of the IS units, each appears to have rod-shaped (α helical) features, though the helices cannot be connected due to limited resolution (Figure S3D). The low visibility of unit X_13_ in the symmetry-released map suggests that this segment is either flexible or occasionally missing; however, at a lower threshold, it looks topologically similar to the others (Figure S3E), and its cross-correlation is thus also similar. Each IS stack is accompanied by a spiral of 13 rod-shaped densities (R_n_) in green (Figure S3E), positioned between the IS components and the tubulin proteins and connecting between each spiral axially in the symmetrized average map (Figure 3G and Figure S3E). Both the number and organization of IS units and rod-shaped densities are also verified in symmetry-released maps (Figure S3E). The unwrapping of the SPMT density map between the protofilaments 4 and 5 further revealed the positioning of IS units with the tubulin lattice from the lumenal side, with unit X_1_ clearly placed over the seam (Figure 3G). Each putative monomeric unit of an IS spans between tubulin heterodimers in adjacent protofilaments and is positioned every 80Å at the interface between β-tubulin and α-tubulin subunits (Figure S4). The IS units marked X_1_-X_13_ are close to β-tubulin at residues 34-42, and the IS units marked R_1_-R_13_ are close to α-tubulin at two loops, residues 30-43 and 359-372 (the second is the S-loop which is longer in α- compared to β-tubulin).

### SPMT seams are oriented non-randomly with respect to the central axis of the parasites

Each IS is coaxial with the tubulin spirals, leaving a gap spanning a portion of the two protofilaments where the seam occurs. The unique arrangement of IS can also be observed in the averaged density map of SPMT segments computationally extracted from the intact parasites (Figure 4A). This can be used to determine the orientation of the microtubule seam. We assigned the 3D SPMT averaged density map back to 5 reconstructed apical complexes of intact parasite cells and retrieved the coordinates applied to align and average the subvolume particles. For each particle, we measured the microtubule seam orientation angle relative to the center of each cell body at cross section, hence the z axis of each SPMT segment. Note that the cell bodies were somewhat flattened, presumably as a result of the preparation and treatments necessary for performing the electron microscopy. For any given tomogram, half the segments showed a preferred orientation >>90°, while the other half had preferred orientations with much smaller angles (<<90°) relative to the z axis (Figure 4B). Overall, we find that the seam orientations’ angle changes according to the position of the SPMT around the cell body and preferentially faces toward the center of the cell body (Figure 4C).

**Figure 4.**
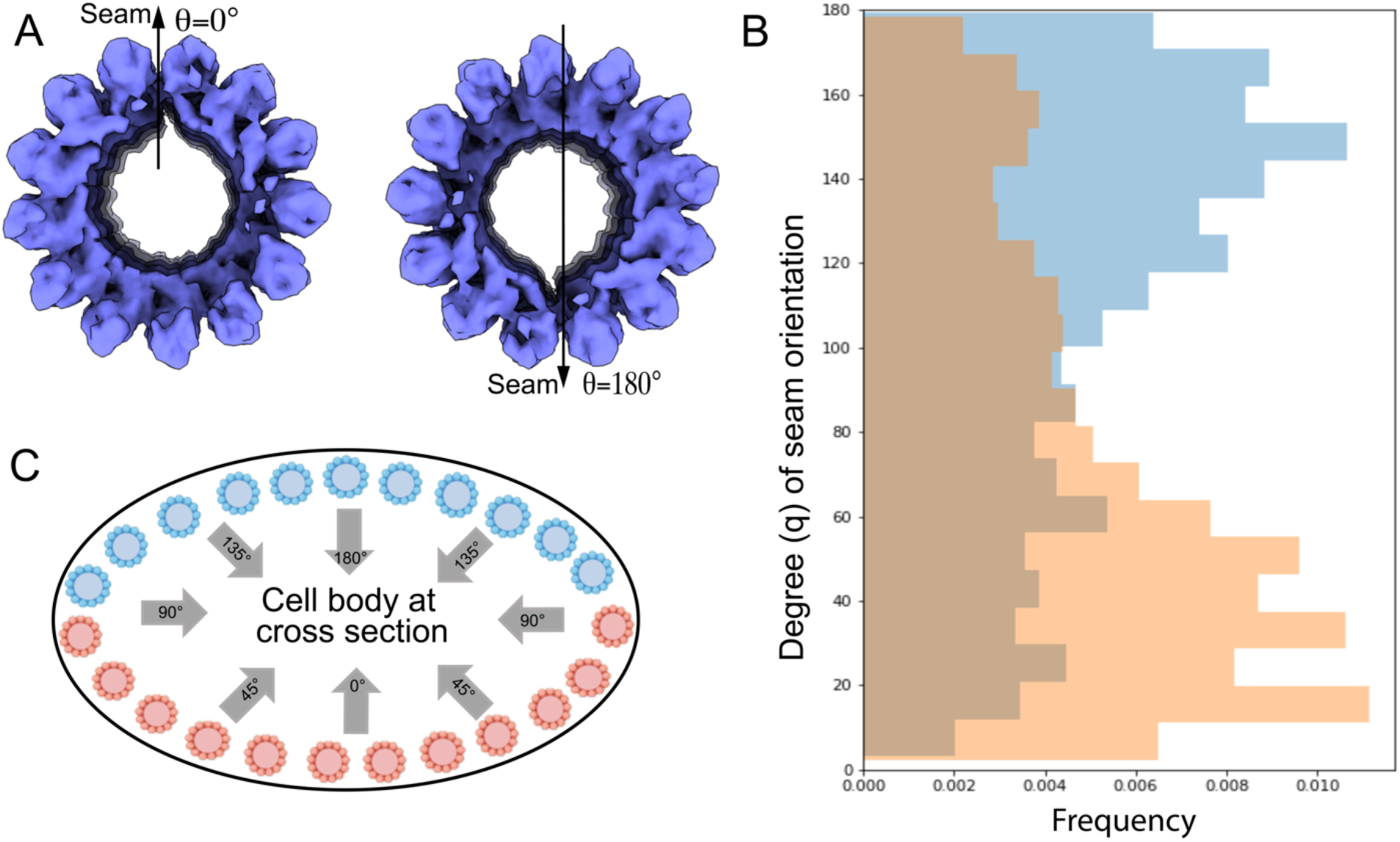
The seam orientation of SPMT in cells is non-random (A) In-cell SPMT segment density is orientated with the seam facing down (θ=180°) or up (θ=0°). We used the gap in the cryo-EM density map of the SPMT segment as a marker of the microtubule seam. (B) The seam of apical microtubules along their axis is analyzed by measuring its angle relative to the z axis in cell tomograms (n=5). (C) A schematic representation of SPMT models in the cell body at cross section, based on the observed cross section of the cell body in the tomograms. The blue color is used for the microtubules in the “top” half layer of the cell (i.e. toward the electron beam), and the orange color is used for the microtubules in the “bottom” half of the cell. Note that the seam is predominantly oriented toward the center of the somewhat flattened cell body, with orientation angle 90° < θ <180° for SPMTs in blue or 0° < θ <90° for SPMTs in orange.

### CFs have a novel tubulin-based structure with additional, unknown densities arranged between the protofilaments

Each CF is tightly associated with its neighbor filament, extending from the top PCR, and forming an array of short, left-handed spirals. Using detergent-extraction to obtain relatively intact cytoskeletons, we were able to refine our subtomogram averaging of the CFs to reveal a periodic density of the CFs (Figure 2B), which was identified to be 8- and 4-nm layer lines in Fourier space, similar to the typical microtubule tubulin repeats (Hu et al., 2002). Given previous work showing that tubulins are a main component of the CFs (Hu et al., 2002; Swedlow et al., 2002), we sought to determine how the tubulins are assembled within these fibrils. To do this, we again used detergent-extracted tachyzoites to obtain their cytoskeleton. This extraction preserves the CFs’ overall helical shape (Figure 2D-H) while allowing the cell body to be much flatter, improving the signal-to-noise ratio for tomographic images. The subtomogram averaged map of the CF was determined at 9.3 Å resolution (Figure S6D). The density map accommodates 9 columns or protofilaments of 40 Å repeated density that we are able to assign as a tubulin subunit using the homology model, together with uncharacterized associated density (Figure 5). Although the data were not of sufficient resolution to visualize the difference between α or β tubulin, published proteomics analysis has shown that the CFs contain both α-tubulin and β tubulin (Hu et al., 2006). Cross-correlation between intra-column units shows the fitting differences between alternating tubulins, suggesting it is polymerized with alternating α- and β-tubulins in a fashion of tail-to-head interactions (Figure S5). The associated components in the center of the CF density map shapes the curvature of 9 tubulin columns with an opening between tubulin column 1 and 9. Three groups of densities are seen within the grooves between column 3 and 4, 5 to 8, and 8 and 9, and these are binding to alternating tubulin units every 80 Å axially (Figure 5B). Compared to the density volume of tubulin monomer (55kDa), the molecular weight of each repeating unit can be estimated to be about 30kDa for each group between column 3 and 4, 95kDa for the ones between column 5 and 8, and 20kDa for the one between column 8 and 9.

**Figure 5.**
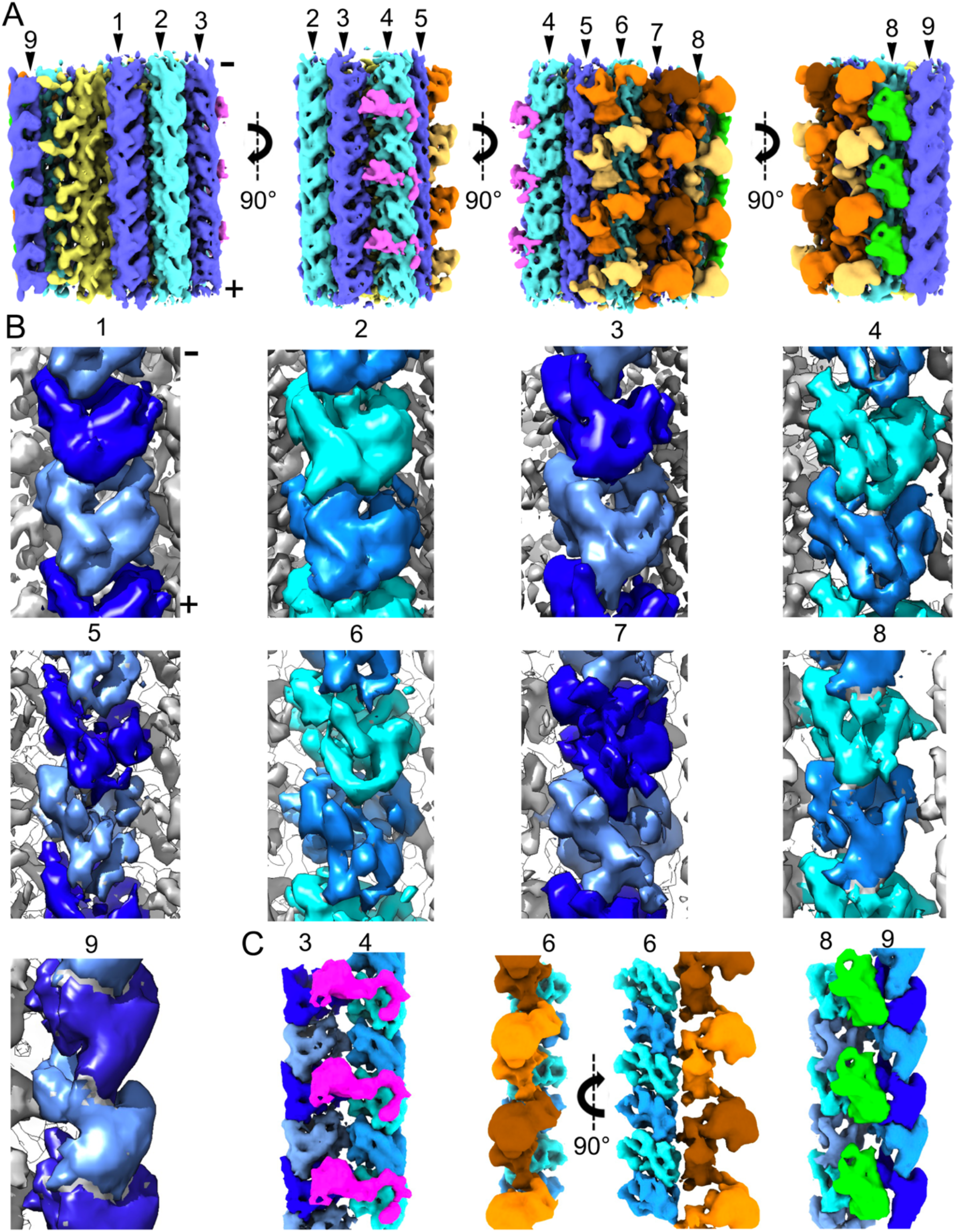
Variable resolvability of tubulin in a CF segment of cryo-EM density (A) CFs are a novel tubulin-based fibril. It comprises 9 columns of tubulin (blue and cyan), associated with CF associated densities within the concave inner face of the CF (yellow) and on the outer, convex face (magenta, green, orange). This is shown axially from various views rotated around the long axis. (B) Zoomed-in cryo-EM density of two tubulin components in each of columns 1-9. Tubulin columns were fitted to the conoid average densities. A total of 9 columns were segmented, shown axially with alternating light and dark blue colors (top) from various views. (C) Non-tubulin groove densities appear every 80Å, shown between columns 3 and 4 (magenta), on the outside of columns 5-8 (orange and brown), and between columns 8 and 9 (green). The plus (+) and minus (−) ends of the microtubule are as indicated in (A) and the first column of (B).

### The plus end of CFs is oriented toward the top of the conoid

With the successful composition assignment of tubulin in the CFs, we were able to determine the structural polarity of the conoid. Figure 5B shows that the plus end of all 9 protofilaments is directed toward the bottom of that figure, based on the fit of tubulin subunits modeled in the density map and by analogy to the plus end of canonical microtubules. This plus direction is toward the top (anterior end) of the conoid. This polarity direction suggests that the conoid is assembled from its base towards what will ultimately be its anterior end. Within the density of extracted CF segments, the 9 tubulin columns are organized into a comma shape in cross-section (Figure 6A). The inter-protofilament angles of CFs are much more variable and often much greater than the relatively constant ~27°of the inter-protofilaments in SPMTs (Figure 6E-G). Interestingly, there is a very pronounced kink between protofilaments 3 and 4, possibly as a result of the observed density between these two protofilaments (magenta in Figure 6).

**Figure 6.**
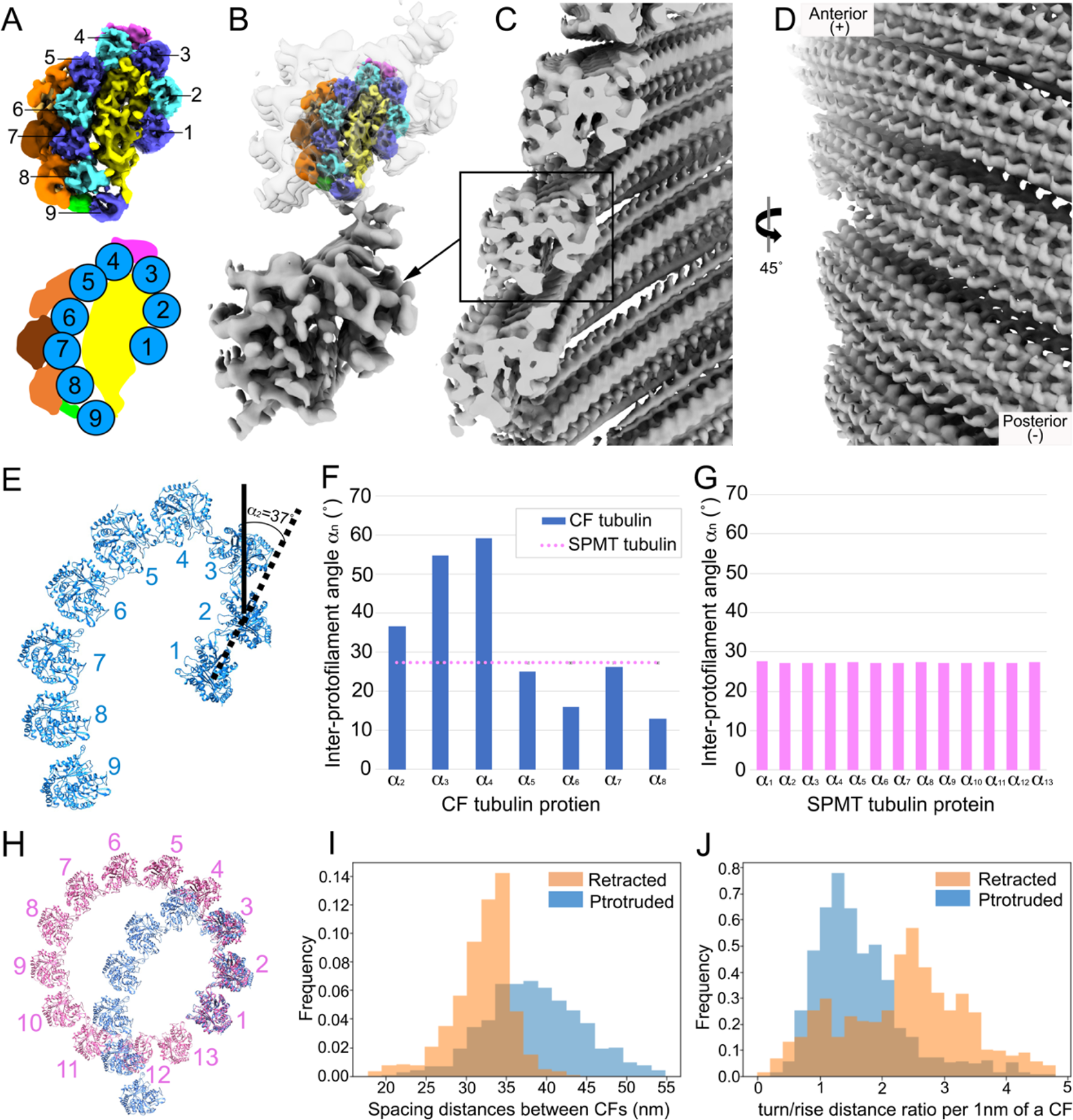
CFs are orientated in parallel with a plus-like end at the anterior (A) Reconstruction of a CF segment from protruded conoids from detergent-extracted cells, generated by subtomogram averaging and shown in cross-section, viewed from the minus-like end. The cartoon (on the bottom) illustrates the structural arrangement of the 9 tubulin columns that form the comma-like architecture. (B) Low resolution reconstruction of a CF segment from conoid-protruded intact cells, generated by subtomogram averaging. Above this is shown the higher resolution structure of part (A) inserted into the low-resolution structure from intact cells. (C) The CFs in a protruded conoid are assembled into spiral filaments with a “parallel” orientation and the opening of the comma-shape facing the interior of the conoid. (D) The spiral CFs viewed from the conoid surface. (E) Zoomed-in model of tubulins within a CF segment. An example of how inter-protofilament angles were determined is shown for the angle between a line connecting the centers of protofilaments 1-2 relative to such a line for protofilaments 2-3; this was defined as α_2_ and was determined to be 37°. (F) The inter-protofilament angles were measured for each protofilament relative to the ones on either side. (G) As for (F) except the inter-protofilament angles were measured for the SPMT protofilaments. (H) Comparison of the protofilament arrangement in the SPMTs and CFs with the tubulin model fitted into the cryo-ET density map. Note the marked kink at protofilaments 3 and 4 in the CF. (I) Closest spacing distance between neighboring CFs showing the increased spacing in protruded CFs (blue) relative to retracted ones (orange). Data are from analysis of 3 protruded and 4 retracted conoids of intact cells. (J) The turn/rise distance ratio was measured every 1 nm along the CF axis showing the CFs in retracted (orange) conoids are higher than in protruded conoids (blue).

Extensive density can be resolved within the structure of the CF segments averaged from intact cells but to a lower resolution than described above for the detergent-extracted cytoskeletons. Nevertheless, we were able to readily fit the higher resolution CF structure into the structure obtained from intact cells with protruded conoids. The results showed considerable extra density within the CFs of intact cells, arrayed along the outside of the tubulin-based protofilaments and associated densities seen for the detergent-extracted material (Figure 6B). Presumably, this extra material dissociates upon detergent extraction. To determine the polarity orientation of the CFs within the cell, their in-situ density was mapped back to the cell tomogram based on the coordinates of each extracted particle. The CF segment density revealed the orientation of the CFs to be in “parallel” with respect to each other, with the opening of the cross-sectional comma shape being toward the center of the conoid (Figure 6C-D). The patterning of retracted and protruded conoids in calcium-ionophore-stimulated parasites was further investigated by measuring the spacing of neighboring fibrils and the ratio between turn and rise distance along the axis under these two conditions (Figure 6I-J). The results indicated that relative to retracted conoids, the distance between neighboring fibrils is slightly increased when the conoid is protruded, with a median spacing of 38.4nm in the protruded conoid vs. 32.6nm in the retracted structures. In addition, the turn-to-rise ratio was markedly lower in the protruded vs. retracted conoids (medians of 1.48 and 2.55, respectively).

## DISCUSSION

Thin or purified samples are ideal to determine near atomic resolution structures of biomolecules in solution using cryo-electron microscopy; however, this usually comes at the expense of studying molecular context in their native environment. Here we developed a protocol for image acquisition and processing by cryo-ET to investigate the cytoskeleton-based apical complex of *Toxoplasma,* examining the cellular machinery in context and extending to the level of single macromolecules. Cryo-ET of intact parasites revealed the spatial organization of distinct components in the highly crowded apical complex. By annotating the tubulin-based SPMTs and CFs, we conclude that their numbers are not fixed, implying that their assembly process is regulated but not rigid (Figures S2D, F). Interestingly, the regulation of number and length of microtubules in *Plasmodium berghei* varied depending on the tubulin expression levels (Spreng et al., 2019). Further investigation into the relationship between SPMT and CF numbers and tubulin expression in *Toxoplasma* will be interesting to study.

The SPMT minus end emanates from the APR, which has been suggested to function like a microtubule organizing center, i.e., as an anchoring site for the SPMTs (Nichols and Chiappino, 1987). Previous studies revealed that even in the absence of fully formed APR, there remains some organization of the SPMT array (Leung et al., 2017). This argued for additional elements that support and stabilize the arrangement of the SPMT. The pillar densities we observe between neighboring SPMTs (Figure 1D) where they attach to the APR could be involved in such organization. The identities of these short densities is not yet known but they may be related to previously described structure found between the SPMTs (Nichols and Chiappino, 1987) and could possibly be comprised of the recently discovered AC9 and/or AC10 proteins which by light microscopy are found just below the APR (Chen et al., 2017a; Tosetti et al., 2020).

How the APR associates with the conoid in parasites with a protruded conoid and whether there is a direct physical connection between the two after conoid protrusion, remains to be determined. RNG2 has been observed to occupy a position consistent with such a role and so could serve to link the two (Katris et al., 2014). In the analyses described here, we did not see any direct connection between these two key structures. There is no evidence to suggest how such a large structure as the conoid can be reversibly protruded if it were physically linked to the APR. On the other hand, we did see intimate connections in the form of discrete filaments connecting the PCR and the anterior ends of the CFs. Enabled by the VPP and subtomogram averaging, we also determined a ratio of 4 PCR subunits for every CF (Figure S1E, F); whether this number differs for conoids with 14 vs. 15 CFs is not yet known and, if it does, whether the number of the PCR subunits determines the number of the CFs (or vice versa), remains to be determined.

Using detergent-extracted parasites and advanced subtomogram averaging protocols for filament structures allowed us to resolve α- and β- tubulin within the SPMTs at subnanometer resolution, as well as observing an inner spiral (IS) of ordered density lining the SPMT lumen as was previously reported but not studied in detail (Cyrklaff et al., 2007; Zabeo et al., 2018). Microtubule assembly by α-tubulin and β- tubulin usually forms a hollow tube, but the formation of microtubules with lumen spirals has been observed in human sperm tails and implicated in flagellar rigidity and directional migration (Zabeo et al., 2018). The presence of IS and the regular orientation of the seam likely play as yet undetermined but critical roles in *Toxoplasma* biology. The SPMTs are reported to be unusually stable in the cold upon ionic detergent treatment (Hu et al., 2002; Morrissette and Sibley, 2002), and their ends are presumably uncapped as seen in sporozoites of the related apicomplexan parasites, *Eimeria* and *Plasmodium* (Cyrklaff et al., 2007; Russell and Burns, 1984), conditions that can normally cause microtubule disassembly (Wallin and Strömberg, 1995; Witman et al., 1972). Assuming the IS complex extends the length of the SPMT, it might serve to stabilize the SPMTs, preventing dynamic turnover from constantly occurring at the plus end. They might also indirectly affect the external surface of the tubulin lattice by inducing some curvature of the microtubule. indeed, the pellicle formed by the SPMTs has a small but distinct helical pitch that correlates with the axis of attachment to solid substrates and the resulting path taken during gliding motility (Frixione et al., 1996; Håkansson et al., 1999; Morrissette et al., 1997). Whether this is inherent to the SPMTs or is an organization mandated by other cellular components, such as the alveolar membranes, is not yet known, but helical reinforcement is an efficient solution regularly used in engineering to provide strength, *e.g*., in carbon-fiber tubes or in armored hoses. The molecular constituents of the IS have yet to be identified but are likely to be among the proteins detected in proteomic studies of purified parasite cytoskeletons (Barylyuk et al., 2020; Gómez de León et al., 2014; Hu et al., 2006). The structural information gleaned here was not of sufficient resolution to allow predictions about such identities to be made but higher resolutions might; alternatively, structural determination of individual, purified proteins would likely enable assignment of the densities to individual molecules in the same way we did here with the well-characterized α- and β- tubulins. Potential mechanisms for controlling the orientation of the SPMT seam, specified by the lumen spirals and which we observed here to have a predictable orientation relative to the cell’s long axis, should be further characterized to understand the role of the seam during motility and invasion.

We were also able to generate a molecular model for the CF tubulin based on the subnanometer cryo-EM density map. By fitting the tubulin subunit model to the density map of the 9 protofilaments comprising a CF, we affirmed that the comma-shaped geometry of the CF is an unusual mode of tubulin polymerization unique to the Apicomplexa. We determined that the assembly of tubulins in the conoid incorporates different tubulin monomers based on the fitting cross-correlation, but we were unable to distinguish between α-tubulin and β-tubulin, as we were able to do for the SPMT, and neither were we able to determine which isotypes of these two proteins were present. The orientation of the tubulin subunits in the CF suggests that it is assembled with a plus end facing the anterior end of the conoid. The densities observed decorating the interior and exterior parts of the tubulin-based portions of the CFs might play a role in the modification of CF curvature. The largest angle (~60°) between protofilaments 3 and 4 seems likely to be related to the groove density uniquely observed in this location (magenta in Figure 6B). Although the identity of the protein(s) represented by this density has not yet been determined, recent studies have shown that TgDCX exclusively localizes to the conoid where it helps stabilize and retain the curvature of the CFs (Leung et al., 2020; Nagayasu et al., 2017) and the size of TgDCX is consistent with the observed density. Also consistent with this possibility, cryo-EM density of conventional microtubules polymerized with bovine brain tubulin *in vitro* shows that the 80 Å periodicity of human DCX is patterned axially between the tubulin protofilaments and that DCX is necessary for the observed curvature of the protofilaments relative to one another in cross-section of microtubules *in vitro* (Bechstedt et al., 2014; Fourniol et al., 2010). TgDCX also induces the curved microtubule arcs in Xenopus S3 cells (Leung et al., 2020).

Conoid protrusion has been suggested to involve modifications in CF patterning (Hu et al., 2002; Leung et al., 2020). The relative angle of each CF to the base of the conoid was revealed to be increased in protruded conoids compared to retracted ones and our data support this observation with a lower turn/rise distance ratio in protruded conoids (Figure 6J), suggesting protruded CFs are less twisted than retracted CFs. We also observed a wider spacing between CFs in the protruded conoid, and it is possible that this is related to the changes in the turn/rise along the fibril length. The progressively decreased resolvability from tubulin protofilaments 1 through 9 in the subtomogram averaging suggests that the “gap” in the comma-shaped cross-section of CFs is flexible, perhaps causing the fibril to twist hence the spacing changes shown in Figure 6I.

Tachyzoites utilize tubulins for different macromolecular assemblies in a novel way. In this study, we substantially refined two molecular structures of the tubulin-based elements in the apical complex, SPMTs and CFs, revealing two very different forms of structural arrangements (Movie S2). This work and its relation to cellular context, therefore, provides new structural detail for the two most substantial forms of tubulin assemblies at the apical end of *Toxoplasma* parasites, the portion of this intracellular parasite that is crucial to the process of host cell invasion. It also provides new understanding of diverse assemblies of macromolecular complexes in the same cell and in those molecules’ native environment.

## STAR METHODS

### Parasite maintenance and cell culture

*Toxoplasma gondii* RH Δ*hxgprt* strain was maintained by growth in confluent primary human foreskin fibroblasts (HFFs) in Dulbecco’s modified Eagle’s medium (DMEM; Invitrogen, Carlsbad, CA) with 10% fetal bovine serum (FBS; HyClone, Logan, UT), 2 mM glutamine, 100 U/ml penicillin, and 100 μg/ml streptomycin (cDMEM) at 37°C in 5% CO_2_.

### Parasite preparation for cryo-EM

HFFs were infected with freshly lysed *Toxoplasma* tachyzoites, and 18-20 hpi, were washed two times with Hank’s balanced salt solution (HBSS without calcium, magnesium and phenol red, Corning, Corning, NY) supplemented with 1 mM magnesium chloride, 1 mM calcium chloride, 10 mM sodium hydrogen carbonate and 20 mM HEPES, pH 7. HFFs were scraped into fresh HBSS and tachyzoites were mechanically released and treated with calcium-ionophore (A23187, Sigma, St. Louis, MO) at a final concentration of 1 μM for 10 minutes at room temperature. Tachyzoites were pelleted and resuspended in fresh HBSS.

For detergent treated tachyzoites, after incubation with calcium ionophore, parasites were pelleted and gently resuspended in 10 mM sodium deoxycholate in PBS, supplemented with protease inhibitor (cOmplete, Sigma, St. Louis, MO). After incubation of 10 minutes at room temperature, tachyzoites were pelleted and immediately loaded on EM grids or resuspended in PBS and pelleted again before loading.

### Cryo-electron Tomography

Plasma treated lacey carbon EM grids were mounted on a manual plunger, loaded with intact or detergent-treated parasite suspension mixed with 10 nm or 6nm gold fiducials (EMS), blotted from the back side using Whatman paper #5, and plunged into ethane cooled down to liquid nitrogen temperature.

Detergent treated tachyzoites were imaged using a Titan Krios electron microscope (Thermo Fisher) equipped with a field emission gun operated at 300kV, an energy filter (Gatan) operated at zero-loss and a K2 Summit direct electron detector (Gatan). High-magnification images of SPMTs and CFs were recorded at 105,000x and 81,000x respectively corresponding to a pixel size 1.38 Å and1.77 Å. Tilt series were recorded using Tomo4 software with bidirectional acquisition schemes (Zheng et al., 2004), each from −45° to 45° with 3° increment. Target defocus was set to −1.0 to −4.0 μm. The K2 camera was operated in dose fractionation mode recording frames every 0.15 or 0.2s. The total dose was limited to 90-100 e/Å^2^ for conoid datasets, and 120-140 e/Å^2^ for SPMT datasets.

The tachyzoites in fresh HBSS were imaged using a Talos Arctica electron microscope (Thermo Fisher) equipped with a field emission gun operated at 200kV, a Volta Phase Plate (Danev et al., 2014), an energy filter (Gatan) operated at zero-loss and a K2 Summit direct electron detector (Gatan). Upon phase plate alignment and conditioning, low-magnification tilt series of intact parasites were recorded at 39,000x at pixel size 3.54 Å using Tomo4 software with bidirectional acquisition schemes, each from −60° to 60° with 2° or 3° increment. Target defocus was set to −1.0 μm. The K2 camera was operated in dose fractionation mode recording frames every 0.6s. Every 2 or 3 tilt series, a new spot on the phase plate was selected. The phase shift spans a range from 0.2-0.8π. The total dose was limited to 70-90 e/Å^2^.

### Tomography Reconstruction and Analysis

The movie frames were motion-corrected using motionCor2 (Zheng et al., 2017), and the resulting micrographs are compiled into tilt series. Tilt series alignment, tomogram reconstruction and CTF estimation are performed automatically using tomography pipeline in EMAN2 (Chen et al., 2019), except the reconstructed tomogram in Figure 1 which was performed by IMOD (Kremer et al., 1996; Mastronarde, 1997). Subcellular features were semi-automatically segmented using the EMAN2 convolutional neural network (Chen et al., 2017b) and refined manually using Chimera (Pettersen et al., 2004).

#### For SPMTs (see flowchart, Figure S6A)

The SPMT from the intact parasite dataset are selected by manual tracing, and subtomograms are generated along the trace. 1,994 particles from 10 tomograms are included in the averaged structure of microtubules from the cellular tomograms. The microtubule pseudo-helical symmetry (Zhang and Nogales, 2015) is applied during the refinement. For SPMT from the detergent-treated dataset, particles are selected automatically using the neural network- based particle picker in EMAN2 (Chen et al., 2019). De novo initial models are directly generated from the particles. The averaged structure of the SPMT is produced from 39,122 particles from 90 tomograms using the subtomogram and subtilt refinement pipeline in EMAN2. The microtubule pseudo-helical symmetry (Zhang and Nogales, 2015) is applied during the refinement, as well as a final symmetry-released map without this symmetry applied to the average.

#### For CFs (see flowchart, Figure S6B)

The CFs from both non-detergent-treated and detergent-treated datasets are selected manually using the filament tracing tool in EMAN2 (Chen et al., 2019). Subtomograms are then generated along the trace with a 75% overlap between neighboring particles. De novo initial models are directly generated from the particles. For the *in situ* CF structure, 1,940 particles from 6 tomograms are used in the subtomogram average. For the CFs from detergent-treated cells, 29,524 particles are generated from 126 tomograms.

#### For both CFs and SPMTs

Subsequent local refinements are performed by extracting sub-particles based on the original subtomogram refinement focusing on different regions of the averaged structure. Further analysis of tubulin-based features, including the orientation of the seam line of microtubules, spacing and curl distribution of the conoid, are computed based on the particle position and orientation from the corresponding subtomogram refinement. Tools for these analyses are distributed within the latest EMAN2 package.

### Map segmentation and modeling

The SPMT map from detergent-treated cells was segmented with Segger (v2.5.4) in Chimera (v1.13), with Grouping by Smoothing, 4 steps of size 1; this generated ~260 regions. Groups of 2 regions appeared to resemble the V-like shape of individual tubulin protein. Chains representing α- and β-tubulin subunits were respectively fitted to groups of 2 regions (with the V-shape) with the SegFit tool in Segger (Pintilie et al., 2010). The fits with the top score matched the map well, with secondary structures in the map and model agreeing well; Z-scores were ~20 (Pintilie and Chiu, 2012), indicating high confidence the fit is correct. The α- and β-subunits were distinguished in the map based on the S-loop which is longer in α- (residues 359-372) vs in β- (residues 358-362) subunits. ISOLDE in ChimeraX was then used to refine the models better into the map (Croll, 2018). The modified models for α- and β-subunits were fitted to each column, creating 13 columns of 6 proteins per column to completely cover the tubulin part of the map.

A map of the symmetrized IS was generated by subtracting densities corresponding to α- and β-tubulin from the entire symmetrized SPMT map. This IS map was then segmented with Segger, without any grouping steps. Rod-shaped regions were grouped interactively, and a cylinder representing an alpha-helix was fitted to each. A beta sheet with 3 strands was fitted to a flat region in the middle. The helices and strands were joined, such that a combined pseudo model consisting of 16 helices and 3 beta strands could be replicated 13 times per spiral, accounting for most observed densities. These units were named X_1_-X_13_. Between the IS and the tubulin protein subunits, 13 further rod-shaped fragments were also observed, and a pseudo model was built for these separately and marked R_1_-R_13_. The helical and beta sheet pseudo-models were built with custom scripts in Chimera. A symmetry released IS map was generated by subtracting densities of α- and β- tubulins from the entire symmetry released map. This map was then segmented with Segger with no grouping steps, and regions were grouped based on which pseudo model of the X or R unit they overlap the most.

Tubulin proteins from SPMT were fitted into the cryo-EM density maps of detergent-extracted CFs by exhaustive search with Situs (Birmanns et al., 2011). Top-scoring fits placed tubulin proteins in columns, much like in the SPMT. Nine columns were found, following a curved open path, with ~8/9 tubulin proteins per column. The fitted proteins were used to mask the CF map (zone tool, 4Å radius) to form the conoid tubulin map. The conoid tubulin map was segmented using the fitted models (using Segger, SegFit, and Group Regions by Chains function). Fitting individual tubulin proteins to each region with Segfit rotational search reproduced the fits, with Z-scores of ~4. Secondary structures in the form of rods are marginally visible in the map, and these corresponded to helices in the fitted models. Extracted tubulin densities from each column were fitted to a single region in the same column using SegFit to study whether there is a repeating α-β-like pattern. The conoid tubulin map was further subtracted from the CF map to form the non-tubulin map. Some repeating densities with clear boundaries could be seen and were extracted with Segger.

## ACKNOWLEDGEMENTS

We thank Dr. Ke Hu and Dr. John M. Murray from Arizona State University for helpful discussion during this work. We thank the support of CZI Biohub Intercampus Team Award for supporting this research. Other support includes NIH grants: S10OD021600, P41GM103832, R01GM079429, P01GM121203, BARD, the United States - Israel Binational Agricultural Research and Development Fund, Vaadia-BARD Postdoctoral Fellowship Award No. FI-582-2018, the Stanford Maternal and Child Health Research Institute, and Stanford School of Medicine Dean’s Postdoctoral Fellowship.

## AUTHOR CONTRIBUTIONS

Conceptualization: W.C. and J.C.B; methodology, M.C., G.P., S.Y.S., L.S.Z.; software, M.C., G.P.; formal analysis, investigation, and visualization, S.Y.S., M.C., G.P., L.S.Z., M.F.S.; resources, S.Y.S., L.S.Z., M.C.; writing – Original Draft, S.Y.S; writing – Review & Editing, S.Y.S., L.S.Z., M.C., G.P., M.F.S., S.J.L., J.C.B., W.C.; funding acquisition and supervision, W.C., J.C.B.

## DATA STATEMENT

The tomograms of intact and detergent extracted *Toxoplasma* apical complex, and subvolume averages of SPMT and CF are deposited to EMDB (accession numbers: xxxx).

