## Supplementary Figures for "Cryo-ET reveals two major tubulin-based cytoskeleton structures in *Toxoplasma gondii*"

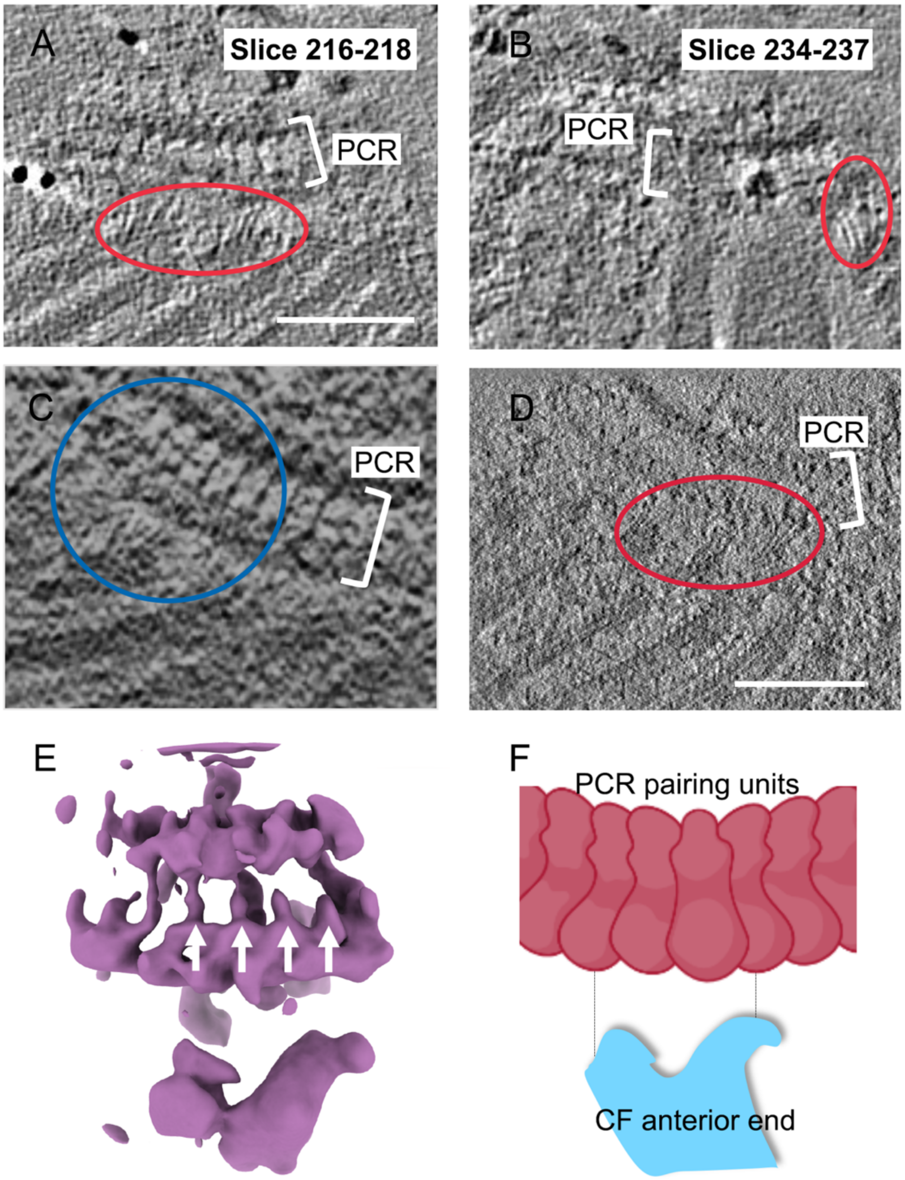


Figure S1. Each CF is associated with 4 PCR units, related to Figure 1
(A-B) Cryo-ET stack showing an intact, non-detergent-extracted parasite with apparent filamentous fibers (circled in red) at anterior end of CFs.
(C) A Cryo-ET slice showing filamentous fibers that connect the CF to the PCR pairing units in an intact cell within a blue circle.
(D) Cryo-ET stack from detergent-extracted CFs and PCRs. Filamentous densities towards the PCR region are highlighted by the red circle.
(E) 3D cryo-EM density map of the blue circle region in (C) by subtomogram averaging. The white arrows point to individual PCR paring units.

(F) schematic of (C) with PCR pairing units (red) and outline of CF anterior end (blue). The filamentous connections occupy the area between them. Scale bar, 100nm.

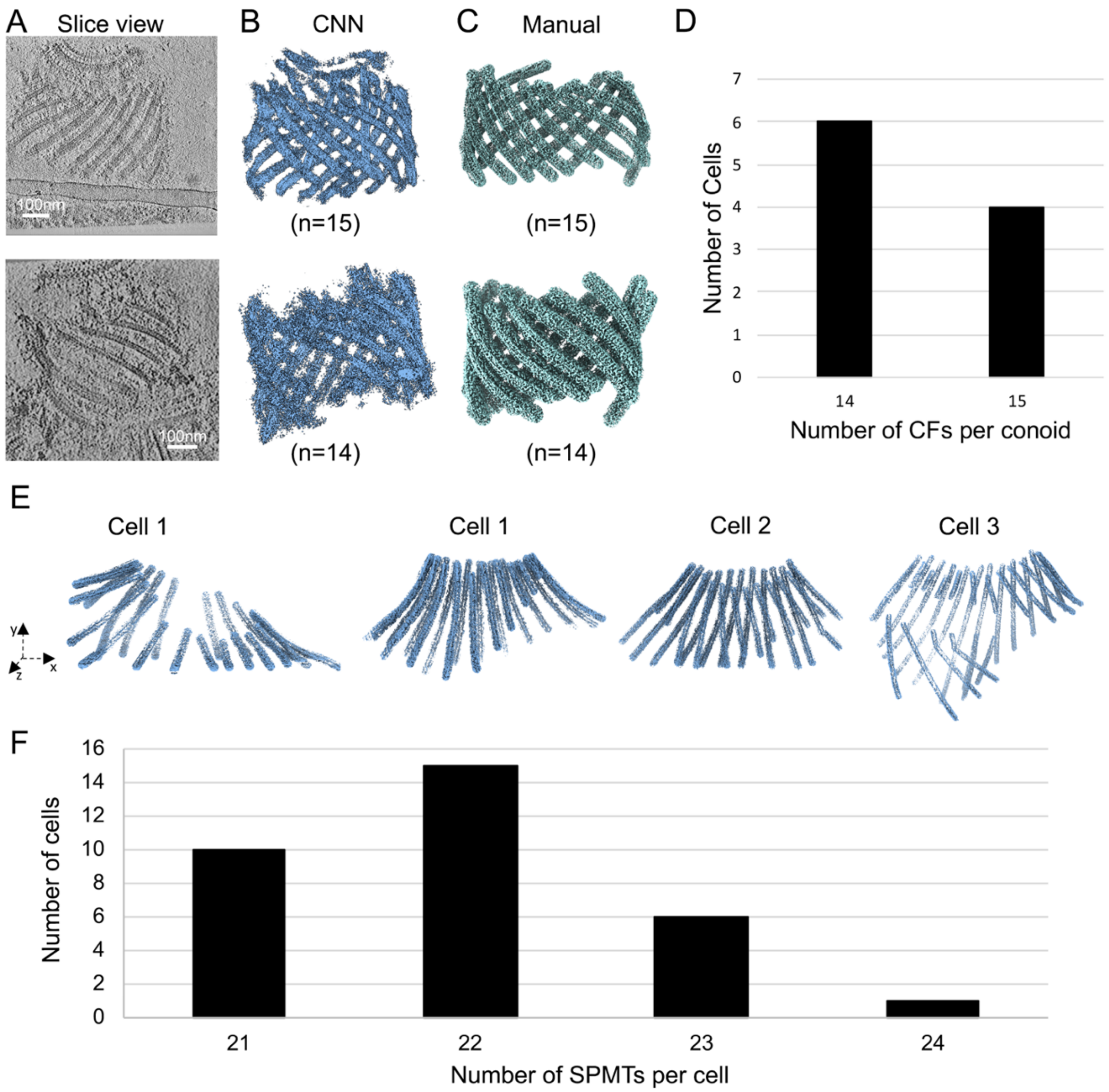


Figure S2. The SPMT and CF number varies between cells, related to Figures 1 and 2
(A) Slice views of two separate tomograms showing the CFs in detergent-extracted cells.
(B) 3D segmentation of the CFs corresponding to the tomograms shown in (A) was performed by CNN.
(C) Manual annotation for CFs. Note that the number of CFs identified in each cell by manual annotation is consistent with the CNN.
(D) The number of CFs per cell is either 14 or 15 (n=10).
(E) 3D segmentation by manually tracing the SPMTs from the APR of representative intact cells showing 21, (cell 1), 22 (cell 2), 23 (cell 3) and 24 (cell 4) SPMTs per cell. Note that only the SPMTs that originate from the APR were counted.
(F) The number of cells with the indicated number of SPMTs per cell counted in 32 different cell tomograms.

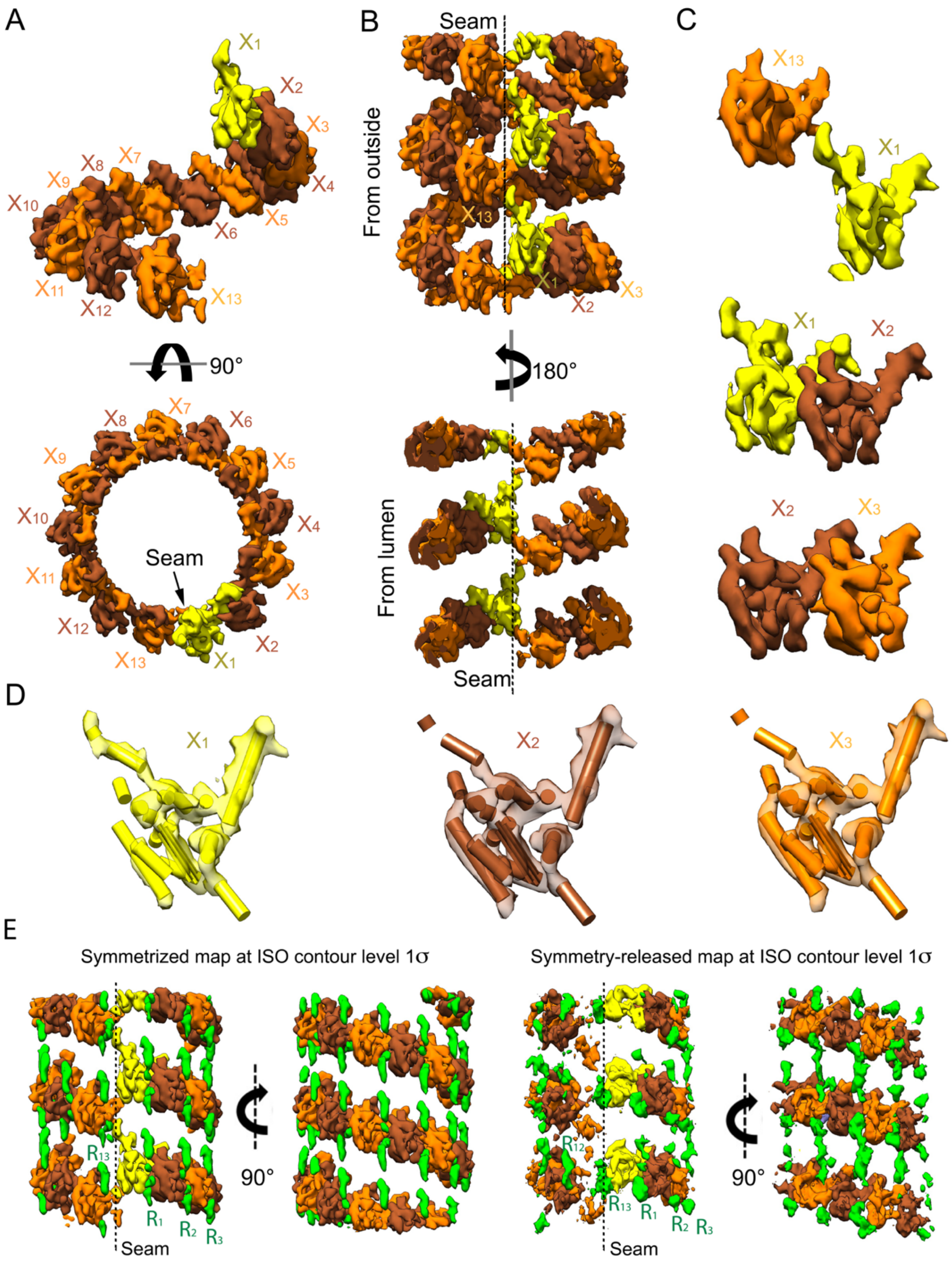


Figure S3. Structural analysis of the inter-IS units, related to Figure 3

(A) One representative stack of IS segmented into 13 units, seen from the seam view (upper panel) and end-on view (lower panel). Note that the vertical distance between the start and end in each stack is 12 nm of helical pitch, and each stack does not extend across the seam.

(B) Multiple stacks of IS segmented into repeating units, viewed from the seam surface (upper) and the lumenal view of the seam (lower). Note each stack has a clear boundary at one end, where unit $X_1$ reaches up toward unit $X_{13}$ at the end of another stack.

(C) Density of consecutive IS units including $X_1$-$X_{13}$, $X_1$-$X_2$ and $X_2$-$X_3$ units.

(D) Representative inter-IS units $X_1$-$X_{13}$ are segmented ($X_1$ - $X_3$ are shown) and a pseudo-model of cylinders and a flat sheet placed into them, based on repeating boundaries of densities.

(E) Representative inter-IS units of symmetrized and symmetry-released SPMT average maps are viewed from seam side or 90˚ rotation at ISO-contour Level 1σ. Note that the unit $X_{13}$ presents low visibility in the symmetry-released map. In addition, the symmetrized maps show the 13 repeating units of an IS stack, accompanied with rod-shaped density $R_1$-$R_{13}$ in green between each IS stack. In the symmetry-released SPMT map, the density of $R_{13}$ is weaker over the seam, similar to the low visibility of $X_{13}$.

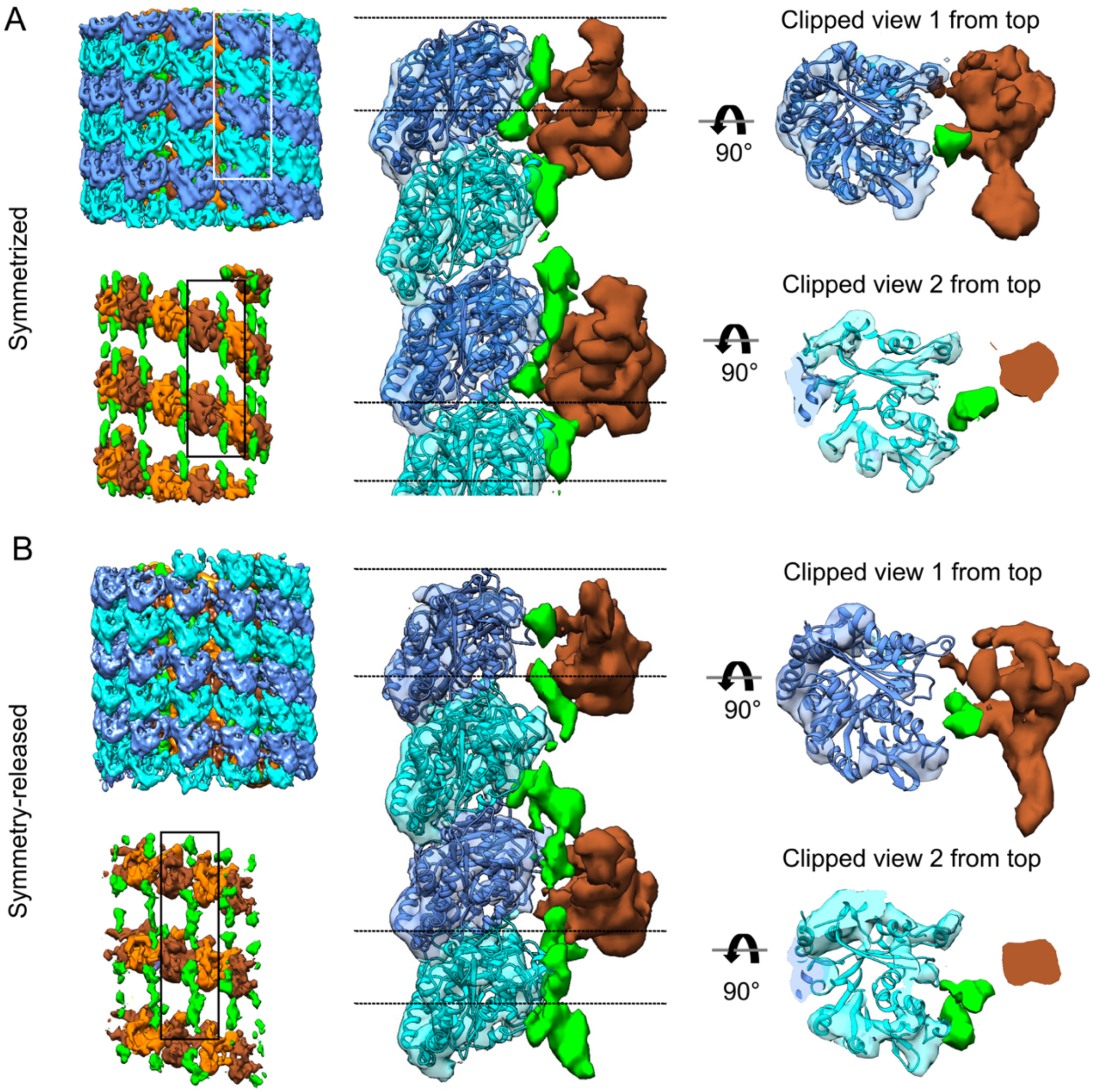


Figure S4. Details of the tubulin-IS interactions, related to Figure 3
(A) The pseudo-helical symmetry-imposed map of the SPMT with the α- (cyan) and β- (blue) tubulin heterodimer segments, viewed from the non-seam side, is shown in the top left. After subtraction of the tubulins' densities, a spiral of likely helical densities (green) associated with each IS stack is apparent along the SPMT axis (bottom left). Middle: a side view of the interaction between the αβ-tubulin and the IS components $X_8$ (sienna) and rod-shaped densities (green). Right: top view and bottom view.
(B) As for (A) except that symmetry-released map is shown (see Methods)

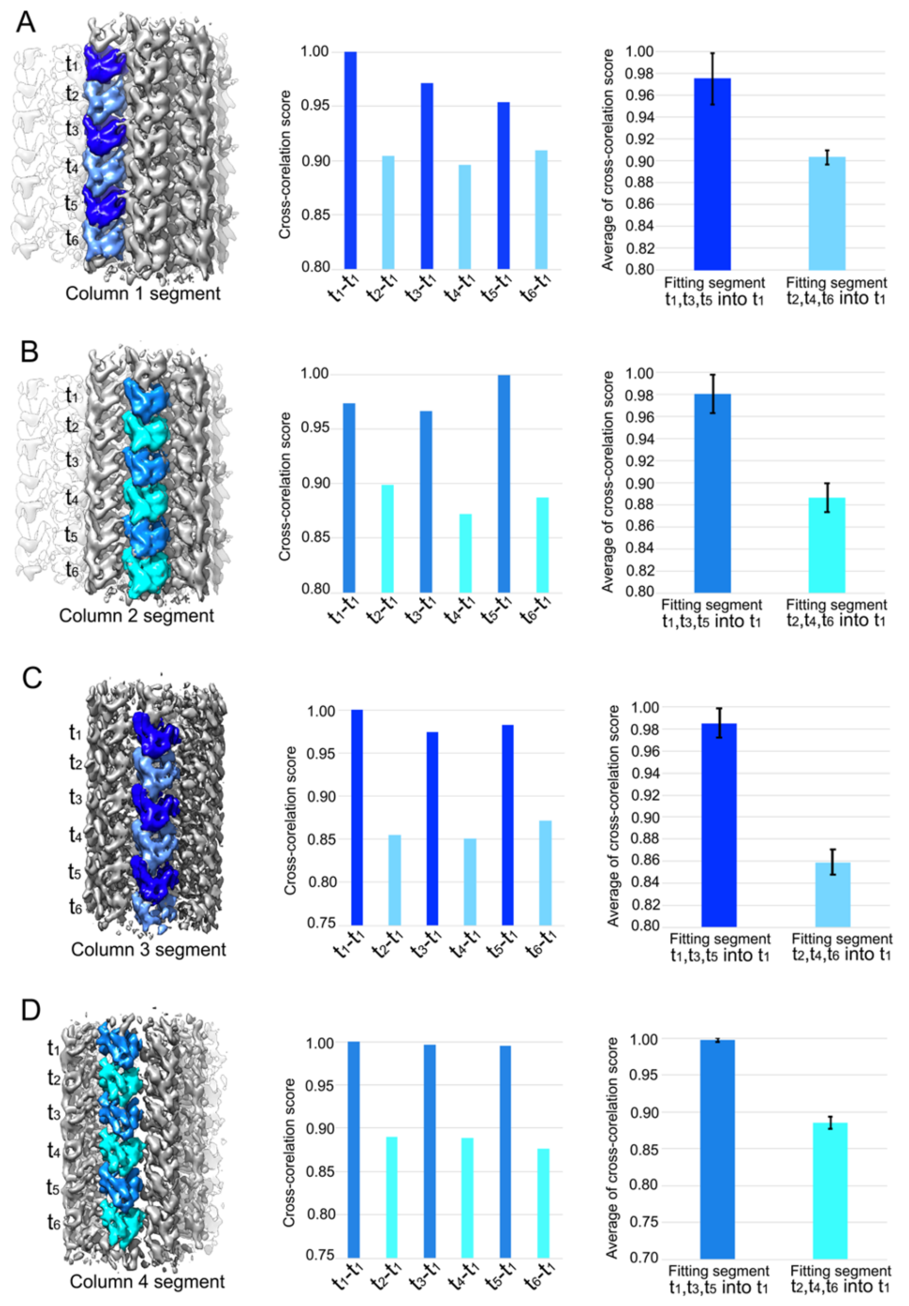
A
B
C
D
t1
t2
t3
t4
t5
t6
Column 1 segment
Column 2 segment
Column 3 segment
Column 4 segment
Cross-corelation score
Average of cross-corelation score
t1-t1
t2-t1
t3-t1
t4-t1
t5-t1
t6-t1
Fitting segment t1,t3,t5 into t1
Fitting segment t2,t4,t6 into t1

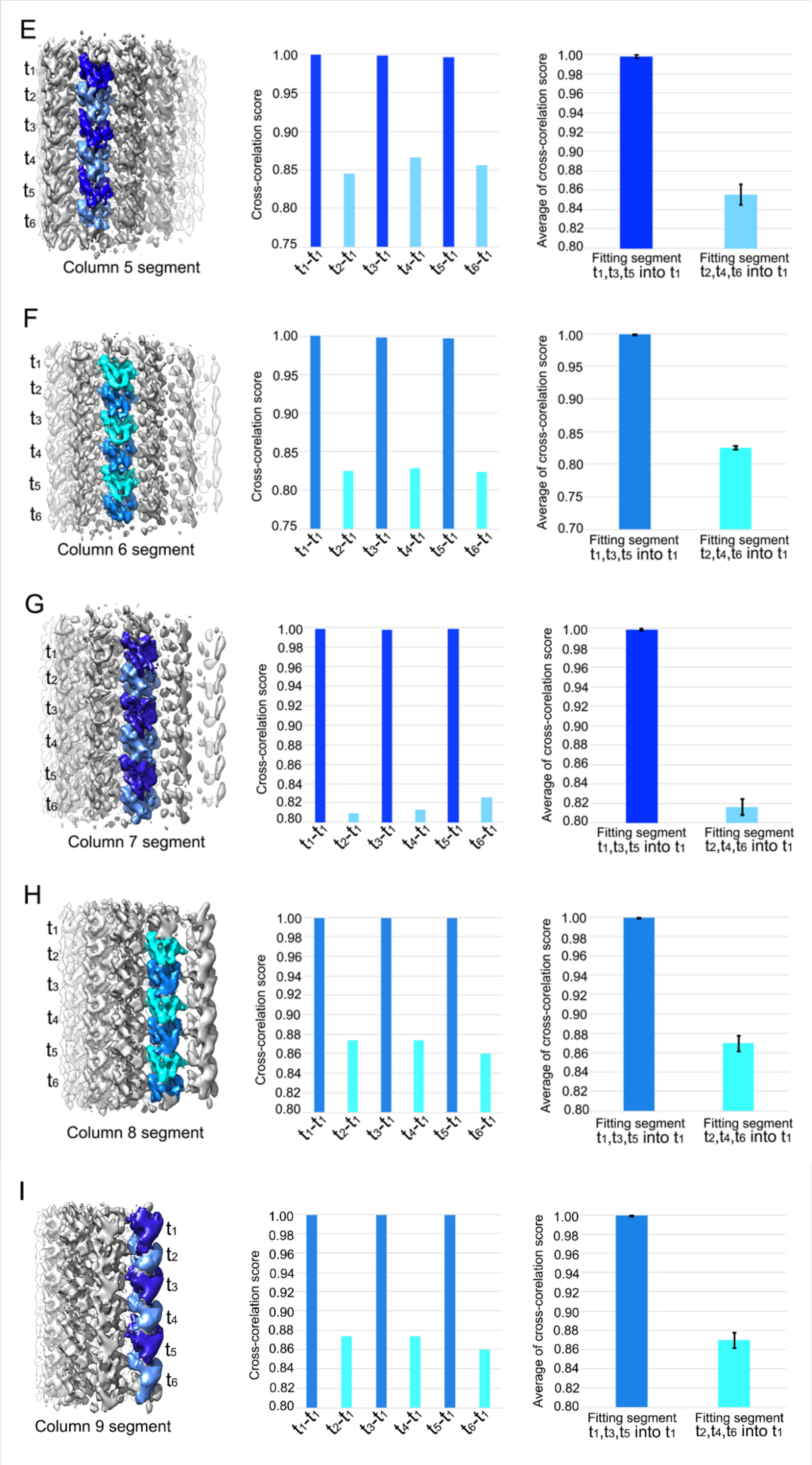


Figure S5. The intra-column comparison for tubulin in the CFs, related to Figure 5

(A) The alternating tubulin in each column (light blue and dark blue) were segmented by fitting the model of tubulin columns into the conoid cryo-EM density map. Each of the 6 segments within a column was fitted to the top segment $t_1$. The cross-correlation score for each fitting is plotted in the middle. Note that the dark blue segments ($t_1$, $t_3$, $t_5$) have higher scores on average than the light blue segments ($t_2$, $t_4$, $t_6$). The averages are plotted on the right, with error bars signifying standard deviation.
(B-I) Scores for columns 2-9 show similar trends, when comparing densities within each column.

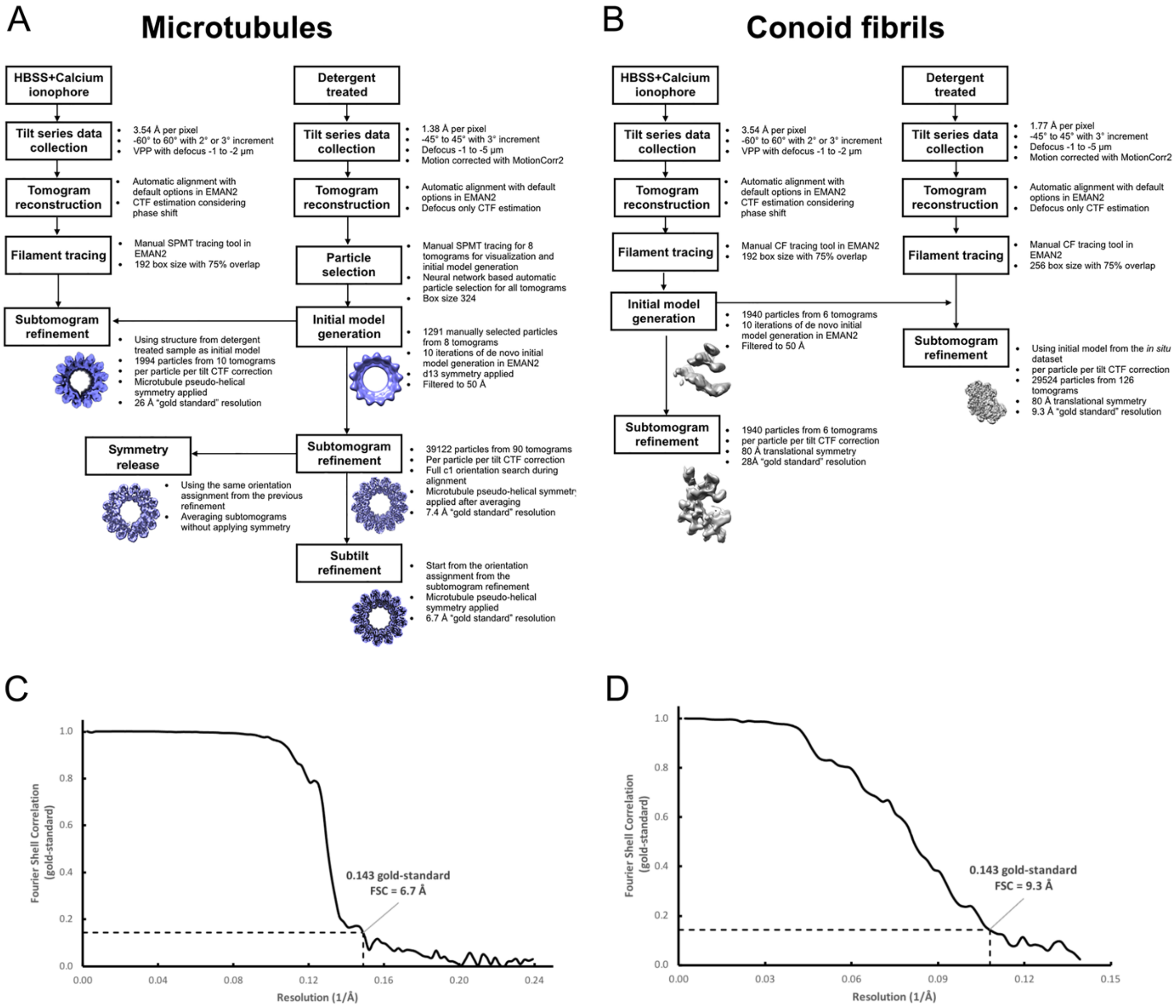


Figure S6. General workflow for subtomogram averaging analysis, related to Figure 2, 3, 4, 5 and 6

(A) The main steps in the workflow of microtubules subtomogram averaging are depicted.

(B) As for (A), the main steps in the workflow of conoid-fibrils are depicted.

(C) Gold-standard FSC curves for refinement for the cryo-EM density map. The resolution of the SPMT map corresponding to the 0.143 FSC cutoff is 6.7Å from two independent sets of subvolume averaged maps.

(D) The resolution of the CF map corresponding to the 0.143 FSC cutoff is 9.3 Å.

**Movie S1**. The movie shows the 3D organization of apical complex in *Toxoplasma gondii* (related to Figure 1).

**Movie S2**. The movie shows the unique structure of two major tubulin-based cytoskeleton in *Toxoplasma gondii* (related to Figure 2,3, 5 and 6).
